# Direct detection of meiotic recombination events in the highly heterozygous amphioxus genome

**DOI:** 10.1101/2025.07.14.664623

**Authors:** Lei Tao, Jing Xue, Junwei Cao, Guang Li, Cai Li

**Affiliations:** State Key Laboratory of Biocontrol, School of Life Sciences, Guangdong Provincial Key Laboratory for Aquatic Economic Animals, Sun Yat-sen University, Guangzhou, Guangdong, China; Guangzhou Institutes of Biomedicine and Health, Chinese Academy of Sciences, Guangzhou, Guangdong, China; School of Life Sciences, Xiamen University, Xiamen, Fujian, China

**Keywords:** amphioxus, recombination, crossover, non-crossover, highly heterozygous genomes

## Abstract

Amphioxus, a basal chordate with highly heterozygous genomes (3.2∼4.2% in sequenced species), represents a key model for understanding vertebrate origins. However, the extreme heterozygosity poses challenges for many genomic analyses, including studying meiotic recombination. Here, we present a novel bioinformatic pipeline that enables direct detection of crossover (CO) and non-crossover (NCO) recombination events using short-read whole-genome sequencing of a two-generation pedigree (two parents and 104 F1 offspring) of the amphioxus *Branchiostoma floridae*. Using parental assemblies generated by Platanus-allee as a custom reference for read alignment, we tracked inheritance patterns in offspring and phased contig-level haplotypes in parents, allowing us to detect recombination events. We identified 2,329 paternal and 2,288 maternal COs, yielding recombination rates of 4.57 cM/Mb and 4.49 cM/Mb, respectively. We found CO coldspots spanning >140 Mb in each parent and these are likely associated with large-scale heterozygous inversions. CO rates were positively correlated with transposable element and gene density in both sexes, but showed weak or no correlation with GC content. We further identified ∼10,000 paternal and ∼5,800 maternal NCO events, predominantly shorter than 200 bp in tract length, and found evidence of GC-biased gene conversion. This work provides the first direct and genome-wide measurement of recombination in amphioxus and demonstrates how high heterozygosity, often considered a barrier, can be leveraged for fine-scale recombination mapping. Our findings illuminate conserved and divergent features of recombination in chordates and establish a framework for studying recombination in other highly heterozygous organisms.

## Introduction

Cephalochordate amphioxus is the sister group of tunicates and vertebrates and represents the most primitive living chordates^[1]^. This key phylogenetic position, together with its simple and slowly evolving morphology and genome structure, makes amphioxus an essential model for studying the origin and evolution of many aspects of vertebrates^[2–6]^.

Genomic research of amphioxus species can offer crucial insights into chordate origin and evolution. The sequenced genomes of amphioxus species span about 500 Mb ^[7]^, with chromosome numbers varying among species (18-20 chromosomes) due to chromosomal rearrangements^[8]^. Notably, amphioxus genomes have a high heterozygosity rate of 3.2-4.2%, significantly higher than humans and most vertebrates, making it one of the most heterozygous known species^[8]^. This high heterozygosity leads to substantial genomic difference among individuals, posing challenges for traditional variant detection methods which are usually based on short-read alignments to a commonly used reference genome. As a result, despite its extensive use in evolutionary studies, population genomic analysis within amphioxus species remains limited.

Meiotic recombination is a fundamental biological process that ensures the proper segregation of homologous chromosomes during gametogenesis and generates novel allele combinations, thereby fueling genetic diversity and facilitating adaptive evolution across eukaryotes^[9]^. Our current understanding of amphioxus recombination rates was based on population genomic variations^[8]^, which reflect historical recombination events averaged over sexes and many generations. As a result, direct experimental observations and quantitative measurements of meiotic recombination— including the absolute number of crossovers per meiosis and their distribution along individual chromosomes—remain completely unexplored in amphioxus.

In this study, we generated short-read whole genome sequencing data for a two-generation family of the amphioxus species *Branchiostoma floridae*, including two parents and 104 F1 offspring. Considering the high heterozygosity of the *B. floridae* genome, we developed a novel bioinformatic pipeline for detecting recombination events in meiosis of the two parents, including crossover (CO) and non-crossover (NCO) gene conversion events. Based on the detected recombination events, we further investigated potential factors influencing recombination in the amphioxus genome. Our work provides novel insights into the recombination landscape of amphioxus and facilitate recombination analysis in species with high-heterozygosity genomes like amphioxus.

## Results

### The workflow for detecting meiotic recombination events

We generated >50X short-read sequencing data for each of the 106 individuals (two parents and 104 offspring). The sequencing depth is not only sufficient for resequencing analysis, but also for high enough for contig-level *de novo* assembly.

Given the high heterozygosity of the amphioxus genome, the existing chromosome-level genome assembly^[7, 8]^ differs significantly from the sequenced genomes in our study. When aligning short reads in our study against the previously used reference genome using short-read aligners, because of low sequence similarity, many reads cannot be correctly mapped, leading to unreliable variant calling results.

To address this issue, we developed a novel strategy for read alignment and variant calling. Because high heterozygosity can facilitate haplotype assembly, we first used Platanus-allee^[10]^, an haplotype assembler designed for highly heterozygous regions, to assemble the parental genomes separately and merged them as a custom reference genome for read alignment of offspring (**Fig. 1a**). As offspring inherit genetic information from parents, most reads from offspring can be correctly mapped to the Platanus-allee-assembled parental genomes, which are though comprised of relatively short contigs. In Platanus-allee, heterozygous regions are assembled as “bubble” contigs with two allelic sequences, named as “primary” and “secondary” respectively. Homozygous regions with identical sequences in an individual are assembled as “non-bubble” contigs (**Fig. 1a**). More details about the assembly and alignment processes are given in a companion paper focusing on *de novo* mutations^[11]^. Based on alignments of offspring reads against parental genomes, we were able to call variants and determine which parental contigs are inherited by each offspring (more details are given in the companion paper^[11]^).

**Fig. 1.**
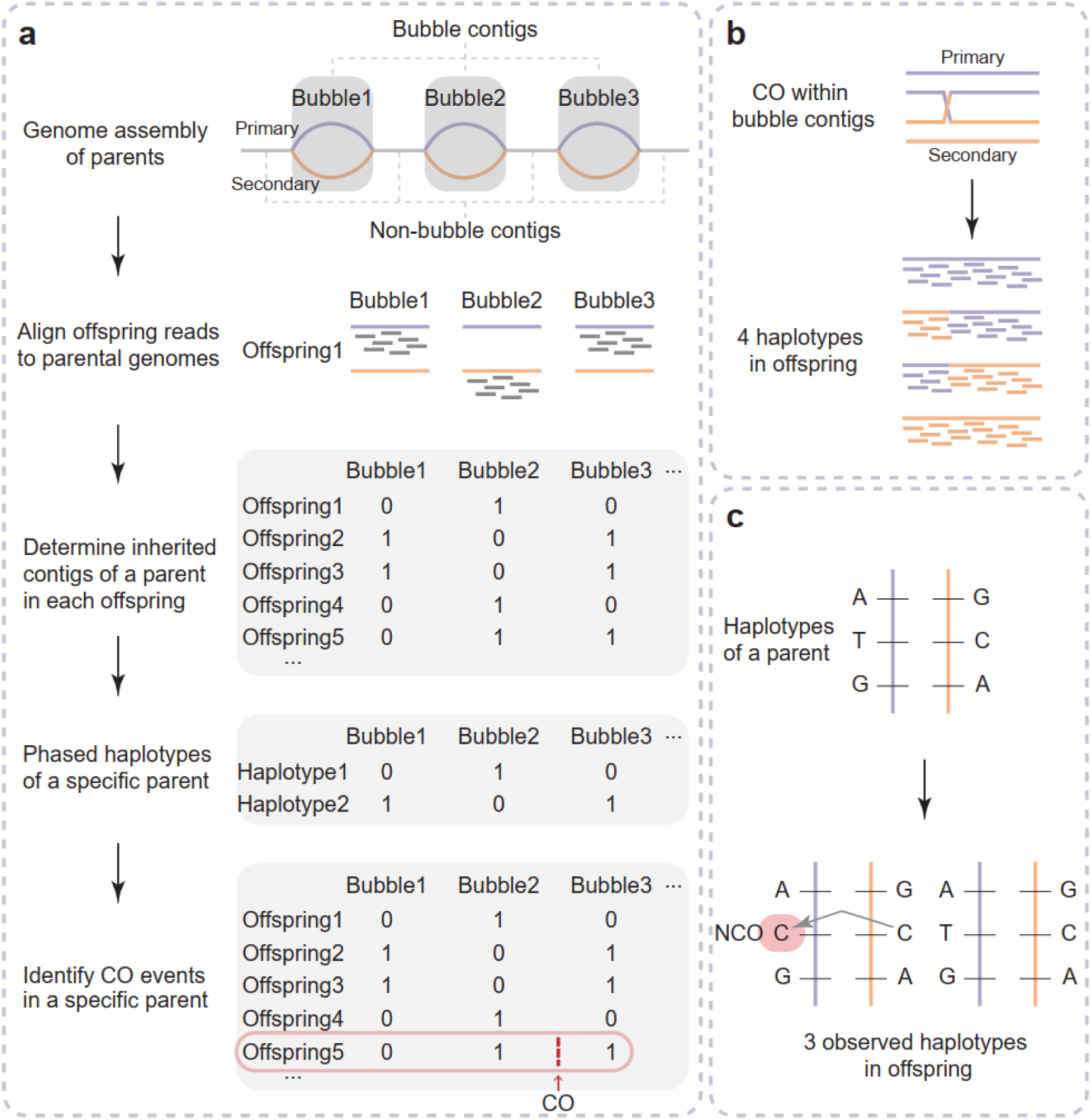
Workflow for detecting recombination events. **a**. Detection workflow for CO events using bubble contigs as markers. By aligning sequencing reads of offspring against the assembled parental genomes, the inherited state of each bubble contig pair of parents in each offspring was determined. Subsequently, the phased contig haplotypes of each parent were determined with contig haplotypes of offspring inherited from that parent. Finally, the CO events in meiosis of a specific parent were identified by comparing parental and offspring haplotypes. The position where an exemplary CO event occurs in Offspring5 is marked with a red arrow. **b**. Detection of CO breakpoints within a bubble contigs pair. A breakpoint caused by a CO event will form four haplotypes in the offspring according to short-reads alignment. **c**. Detection workflow for NCO events. Within a pair of bubble contigs, a portion of one contig replaces the allelic region of the other, giving rise to three observable haplotypes in offspring.

Since non-bubble regions in the parents have identical haplotypes across all offspring and are not informative for recombination analysis, we thus only focused on bubble contigs. The paternal genome assembly has 42,378 bubble contig pairs (∼466.3 Mb in total, average contig length of ∼5.5 kb), and the maternal one has 43,292 pairs (∼389.9 Mb in total, average contig length of ∼4.5 kb). For bubble regions of a specific parent, each offspring can only inherit either the primary or secondary contig, so such regions can be considered bi-allele markers for phasing analysis. Using hapi^[12]^ with offspring states of bubble contigs as markers, we performed parent-level phasing of bubble contigs and reconstructed parental haplotypes for each parent (**Fig. 1a**). Our method is analogous to that of using gamete sequencing data to reconstruct parental haplotypes^[13, 14]^, though our method has much easier sample preparation and affordable sequencing cost. After phasing, by comparing offspring haplotypes and reconstructed haplotypes from a specific parent, we can detect CO recombination events occurring in the meiosis of that parent (**Fig. 1a**).

Unlike CO detection based on single nucleotide polymorphism (SNP) markers, bubble contigs can have internal COs, especially for longer contigs. Therefore, we also analyzed putative CO-induced breakpoints within contigs based the read alignment of offspring reads against parental bubble contigs (**Fig. 1b**). After recombination, related sequences from a bubble contig pair form four possible haplotypes in the offspring. Though the number of breakpoints that can be found within contigs is limited, this analysis can complement the CO detection method that used bubble contigs as markers. In addition to CO events, we also detected NCO gene conversion events in the genome. NCO events usually have a smaller impact sequence range. For instance, the average tract length of NCO events in humans is 459 bp^[15]^. Due to few informative markers, we cannot identify NCO events in non-bubble regions, so we focused on those in bubble regions. To detect NCO events, we compared the primary and secondary contig in each bubble contig pair and identified high-quality heterozygous variants as informative markers. An NCO event was considered to have occurred when the parental phase of first and last markers on a contig remained unchanged while the middle marker(s) switched to another allelic type (**Fig. 1c**). For NCO events, we also required that only three haplotypes were observed in the offspring.

### Characterization of CO events

Based on the inheritance patterns of bubble contigs in offspring samples, we reconstructed parental haplotypes for each parent, using bubble contigs as markers. This allowed us to identify CO events during meiosis in the father and the mother, respectively. In total, we identified 2,329 CO events from paternal meioses across all offspring, averaging 22.4 per meiosis (range: 13-37) (**Fig. 2a, Supplementary Table 1**). For maternal meioses, we identified 2,288 CO events, averaging 22.0 per meiosis (range: 10-37) (**Fig. 2a, Supplementary Table 1**). In most diploid sexually reproducing species, there is usually at least one CO event per meiosis on average^[16–18]^. The CO frequency in this amphioxus family aligns well with this pattern. As the haploid genome of *B. floridae* is about 490.4 Mb^[8]^, the CO recombination rate is 4.57 cM/Mb in the father and 4.49 cM/Mb in the mother, higher than that in mammals (∼0.5 cM/Mb) and birds (∼2 cM/Mb)^[19, 20]^. Unlike in humans and some other vertebrates where a certain specific sex has higher CO rates^[21, 22]^, we found no significant sex biased CO rate in this *B. floridae* family.

**Fig. 2.**
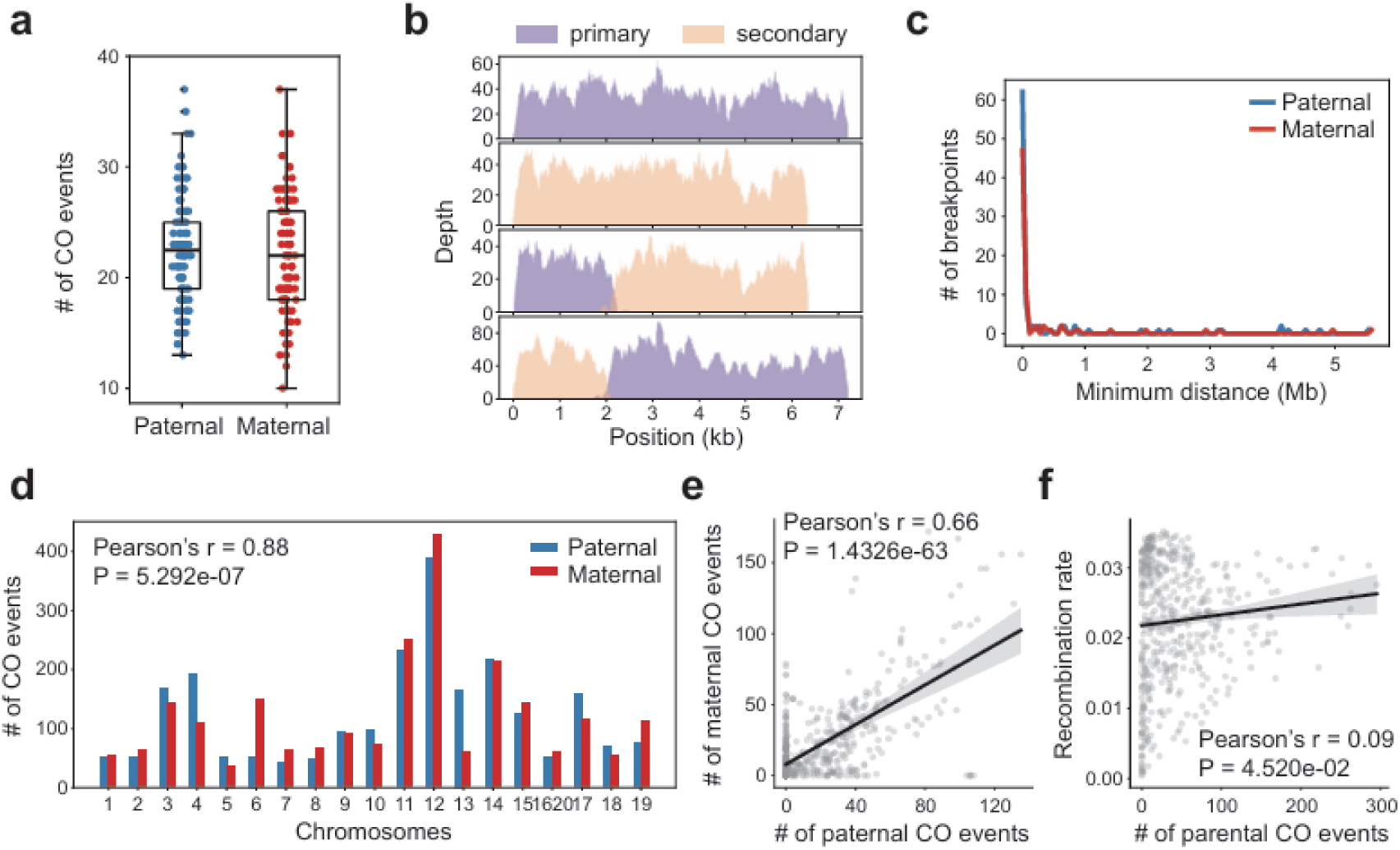
Characterization of CO events in paternal and maternal genomes. **a**. Numbers of CO events observed in 104 offspring, separating paternal and maternal COs. **b**. An example of a CO breakpoint within a bubble contigs pair. Sequencing coverage depth plots are shown for four offspring aligned to the bubble contigs region. From top to bottom, these four samples respectively inherited the primary contig, the secondary contig, a new contig formed by the front segment of the primary contig combined with the rear segment of the secondary contig, and another new contig formed by the rear segment of the primary contig combined with the front segment of the secondary contig, representing 4 distinct haplotypes. **c**. Distribution of distances between all identified within-contig CO breakpoints and their nearest detected CO events using bubble contigs as markers. **d**. Numbers of identified CO events per chromosome in the father and the mother. **e**. Correlation between the genomic distributions of CO events identified in the father and those in the mother. The window size was set to 5 Mb, with a step size of 1 Mb. Linear regression was employed to fit the data, with the 95% confidence interval as shaded area. **f**. Correlation between the distribution of CO events and recombination rates estimated by Huang et al.^[8]^ The window size was set to 5 Mb, with a step size of 1 Mb. Linear regression was employed to fit the data, with the 95% confidence interval as shaded area.

We further identified putative CO breakpoints within bubble contigs. These recombined regions swapped parts of two paired bubble contigs, creating four haplotypes in the offspring (**Fig. 2b**). We identified 171 putative breakpoints (95 paternal and 76 maternal). Most breakpoints (∼90.6%) were within 1 Mb of the nearest CO event detected by the bubble contig inheritance method (**Fig. 2c**), suggesting they represent the same CO events. This also implies that our strategy using bubble contigs as markers effectively detected most genomic CO events.

Subsequently, we analyzed the distribution of CO events across different chromosomes. We observed a higher number and density of CO events on certain chromosomes, such as chromosomes 11, 12, and 14 (**Fig. 2d**). This may relate to specific genomic characteristics, a topic we will discuss later. Most chromosomes showed limited sexual dimorphism in the number of CO events (**Fig. 2d**). Additionally, the distribution of CO events on chromosomes in the father and the mother exhibited a high correlation (**Fig. 2e**, Pearson’s r = 0.66, in windows of 5 Mb). This suggests that in this amphioxus pedigree, sex difference has limited impact on both the quantity and chromosomal distribution of CO events. We also compared our CO results with the population-level CO rates estimated by Huang et al.^[8]^ based on population genomic variations. The correlation between our pedigree-based CO rates and previous population-level CO rates is weak (Pearson’s r = 0.09, p = 0.045, in windows of 5 Mb) (**Fig. 2f**), likely due to the fact that the two methods focus on different CO events. Huang et al. focused on the historical CO events at the population level which can be affected by evolutionary processes such as selection, but our data focus on contemporary CO events derived from a single family.

A sex-determining region of *B. floridae* spanning 0–3.1 Mb of chromosome 1620 (resulting from a chr16-chr20 fusion) was reported previously^[8]^. This species exhibits a ZW sex determination system, with males being homogametic (ZZ) and females heterogametic (ZW)^[23]^. In the analyzed pedigree, CO events were detected near the sex-determining region in the paternal genome but absent in the maternal genome, consistent with suppressed recombination in the heterogametic sex.

### Impact of heterozygous inversions on the distribution of CO events

When analyzing CO distribution across the genome, we noticed that although the overall correlation of CO events on chromosomes in the father and the mother was relatively high, there are large genomic regions without any CO events, which differ among paternal and maternal genomes (**Fig. 3a**). For each parent, we defined regions spanning 5 Mb or more with no CO events in any offspring as coldspot regions. The paternal genome had ∼209.1 Mb of CO coldspot regions (genomic coordinates in **Supplementary Table 2**), and the maternal genome had ∼141.9 Mb (**Supplementary Table 2**). One of the reasons for the formation of coldspot regions might be the inhibitory effect on peripheral CO events near the centromere^[24, 25]^, but this cannot explain the existence of coldspot regions at positions far from the centromere. Another point that easily comes to mind is that the sparsity of markers (bubble contigs) in these regions leads to a decrease in the sensitivity of detecting CO events. However, in many cases, coldspot regions had high marker densities, and COs were also detected in marker-sparse regions. For example, on chromosome 3 of the maternal genome, CO events were mainly observed in the marker-sparse half (16.9-32.3 Mb), while the marker-dense half (0-16.9 Mb) had few COs (**Fig. 3a**). This suggests that coldspot regions are not simply due to marker sparsity.

**Fig. 3.**
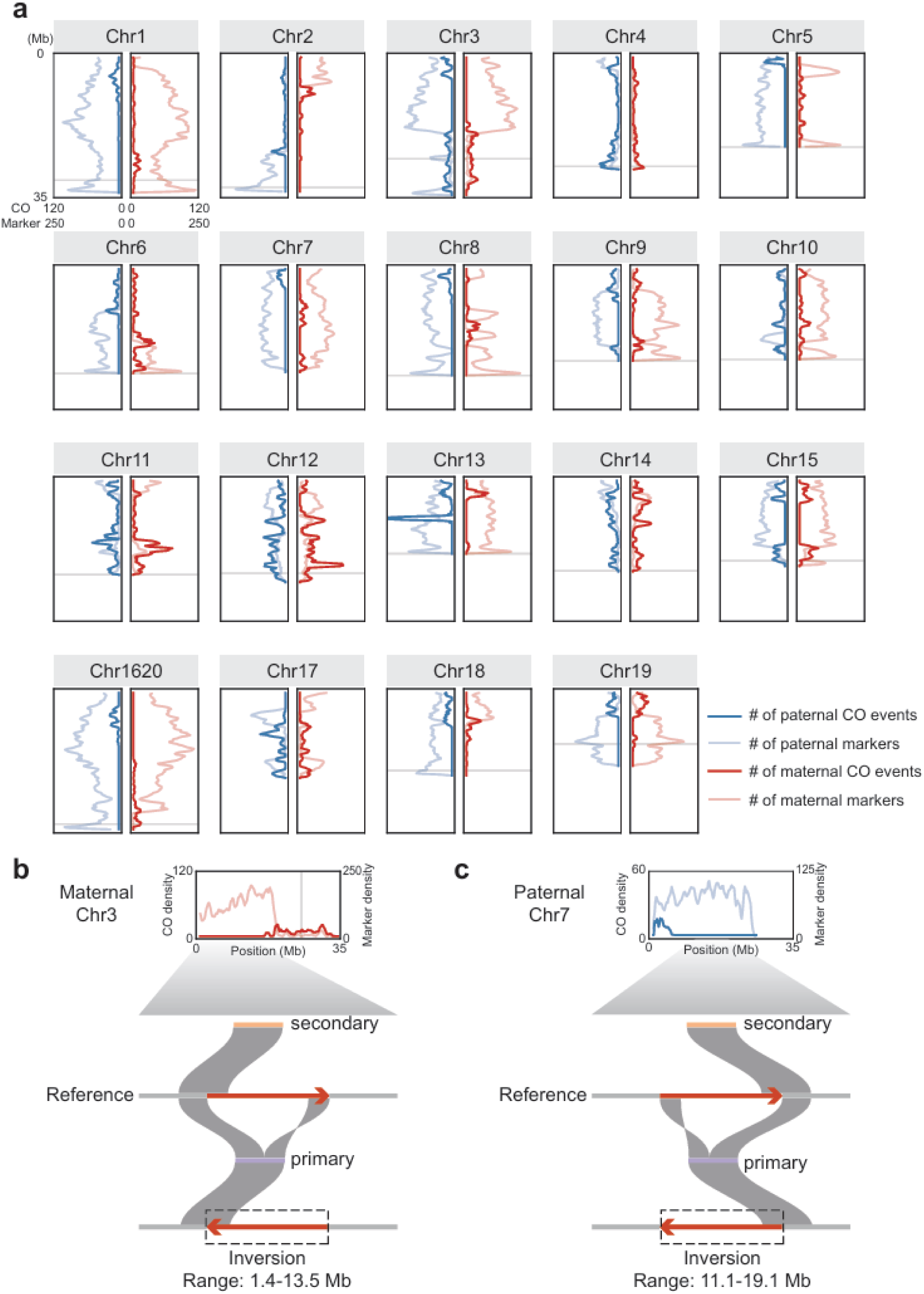
Distribution of COs on chromosomes and examples of large inversions potentially suppressing COs. **a**. Densities of identified CO events and bubble contig markers along the paternal and maternal genomes, respectively. A window size of 1 Mb was used. Centromeric regions (genomic coordinates from Huang et al.^[8]^) are shaded. Centromere positions for chromosomes 7 and 17 were not definitively identified. (**b**) and (**c**) respectively illustrate examples of heterozygous inversions in the maternal and paternal genomes. Shown are pairwise alignments of primary and secondary contigs from one set of bubble contigs against the chromosome-level reference genome assembly^[8]^.

Previous studies suggest that large-scale structural variations, especially heterozygous inversions, can suppress CO events^[26–28]^. The amphioxus genome exhibits substantially elevated rates of structural variations, with the rate of translocations and inversions reported to be approximately 30 times of that observed in the human-chimpanzee comparison^[29]^. To check if such variations cause CO coldspots in amphioxus meiosis, we searched for large-scale inversions in parents using phased haplotype information. We found four large-scale heterozygous inversions in the parental genomes and all of them are located within CO coldspots (**Supplementary Table 3**). For example, around the position of 1.4 Mb on maternal chromosome 3, the secondary contig of a bubble contig pair aligns with this region nicely, but the corresponding primary contig can be mapped to this region only partially, with the other half being inversely aligned to a region around 13.5 Mb on the same chromosome (**Fig. 3b**). This implies a large-scale inversion (involving 1.4 Mb to 13.5 Mb of chromosome 3) in the primary contig-carrying haplotype, likely suppressing COs during maternal meiosis and forming a maternal CO coldspot. A similar case was found in chromosome 7 of the paternal genome (**Fig. 3c**). However, due to the limitations of genome assemblies from short read sequencing data, we cannot confirm whether all coldspot regions harbor large heterozygous inversions. Nonetheless, we propose that such heterozygous inversions are likely one main cause for CO coldspots in the amphioxus genome.

### Association between CO events and genomic features

We further investigated the relationship between CO rates and genomic features, including transposable element (TE) density, gene density, and GC content^[30]^. In both paternal and maternal genomes, TE density showed significant positive correlations with CO events (Pearson’s r of 0.43 in father and 0.40 in mother, **Fig. 4a**), suggesting that TEs are a likely factor influencing COs in amphioxus. Further analysis of different TE types showed that SINEs, LINEs, LTRs, MITEs and DNA transposons all correlate with CO distribution respectively (**Fig. 4b**), without a clear preference for any specific TE class. Many previous studies found a negative correlation between TE density and CO rate^[31, 32]^, though some examples of positive correlation were also reported^[33–35]^, implying the complex relationship between TEs and CO events. Gene density also correlated with CO events (Pearson’s r of 0.28 in father and 0.26 in mother, **Fig. 4c**), but less strongly than TE density. Regions with high gene density tend to have higher CO rates, possibly due to more accessible chromatin structure in gene-dense regions^[36]^, which facilitates the binding of recombination-associated proteins and promotes recombination. No significant correlation was found between GC content and paternal COs, while a weak negative correlation was observed in maternal genome (Pearson’s r = −0.12, p-value = 0.0164, **Fig. 4d**). Overall, paternal and maternal genomes showed similar patterns: higher TE and gene densities tend to be associated with higher CO rates.

**Fig. 4.**
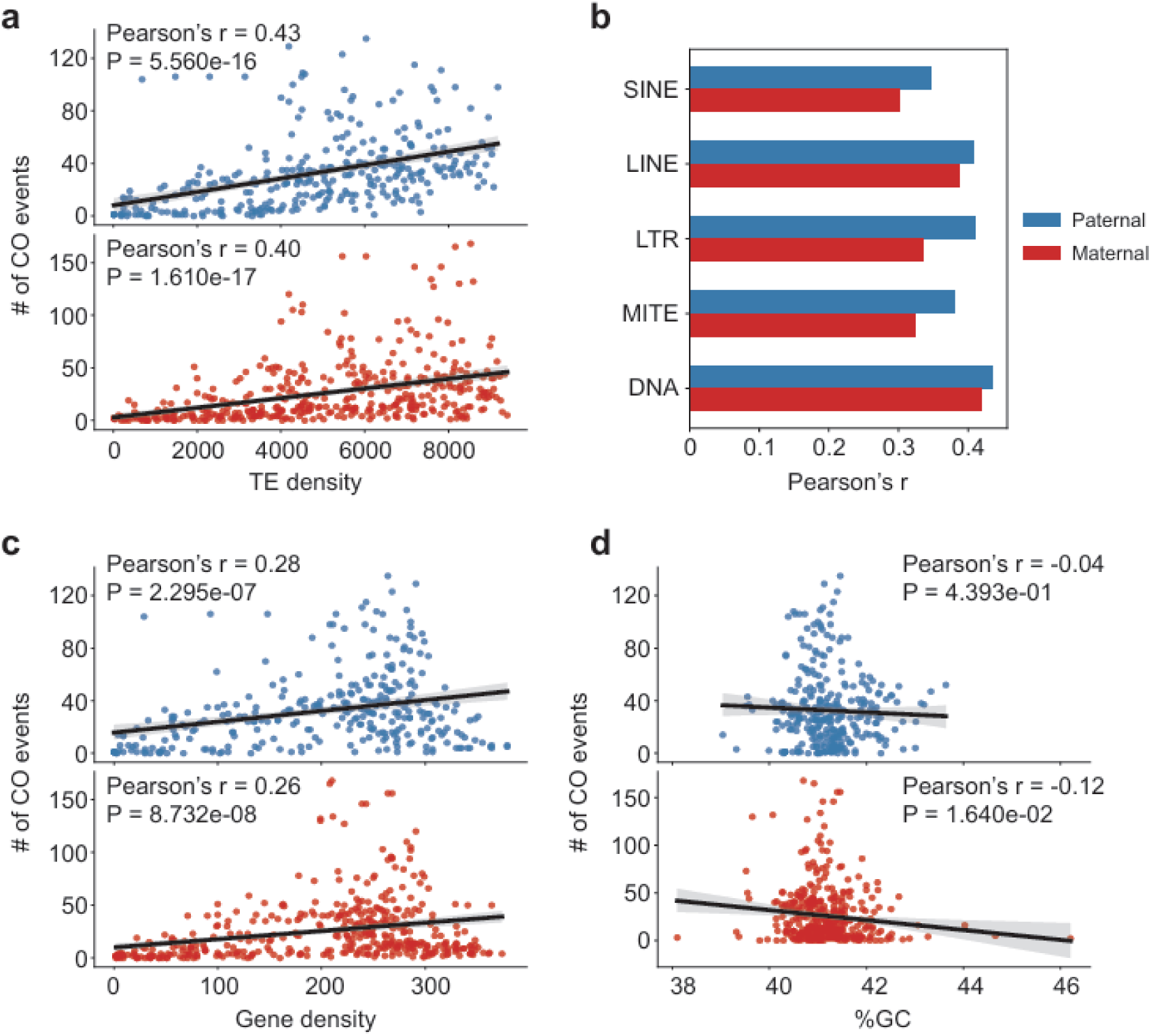
Association between CO events and genomic features. **a**. Scatter plots displaying the relationship between CO density and TE density, with Pearson’s correlation coefficients in the figures. The window size for calculating densities was set to 5 Mb, with a step size of 1 Mb. The blue scatter plot at the top represents paternal data, and the red scatter plot at the bottom represents maternal data. Linear regression was employed to fit the data, with the 95% confidence interval as shaded area. **b**. Pearson’s correlation coefficients between CO density and the densities of specific TE types. All correlations shown have p-values < 0.001. **c**. Scatter plots displaying the relationship between CO density and gene density. Window size, dot color and data fitting are the same as in panel a. **d**. Scatter plots displaying the relationship between CO density and GC content. Window size, dot color and data fitting are the same as in panel a.

### Characterization of NCO events

Apart from CO events, recombination can generate NCO gene conversion events. As NCO tract lengths are usually short, detecting NCO events is highly dependent on the density of informative heterozygous SNP markers surrounding NCOs. The high heterozygosity of *B. floridae* individuals provides a unique advantage for identifying NCOs. Due to insufficient SNP markers in non-bubble regions and some bubble regions with few differences between primary and secondary contigs, we first identified the genomic regions where NCO events could be detected (see Methods). In the paternal genome, an average of 73.0 Mb of the genome for each offspring was analyzable for NCO events, with 9,992 NCOs detected (63-141 per offspring, averaging 96.1, **Fig. 5a, Supplementary Table 4**). In the maternal genome, an average of 61.1 Mb for each offspring was analyzable, with 5,850 NCOs found (25-114 per offspring, averaging 56.3, **Fig. 5a, Supplementary Table 4**). Overall, more NCO events were observed in the paternal genome. However, unlike CO event detection with bubble contigs, NCO detection is more affected by marker density. Thus, it is unclear whether the higher NCO count in the paternal meiosis reflects a true biological difference like humans^[37]^ or is due to more informative paternal markers (6.71 million versus 5.78 million markers). Assuming the analyzable regions are representative of the whole genome, the genome wide NCO:CO ratios are 28.8 and 20.5 for paternal and maternal meiosis respectively. These ratios in humans are 7.84 in male and 3.91 in female^[37]^. It is also higher than some other vertebrates, such as zebra finch (5.4)^[38]^. The high ratios in amphioxus could be due to the amphioxus genome’s high heterozygosity that enhances NCO detection sensitivity. We note that although it is impossible to distinguish an NCO event from a *de novo* substitution at the parental heterozygous sites with our method, given the low mutation rate^[11]^, the level of NCO false positives due to *de novo* substitutions at heterozygous sites should be very low (more information given in the Methods).

**Fig. 5.**
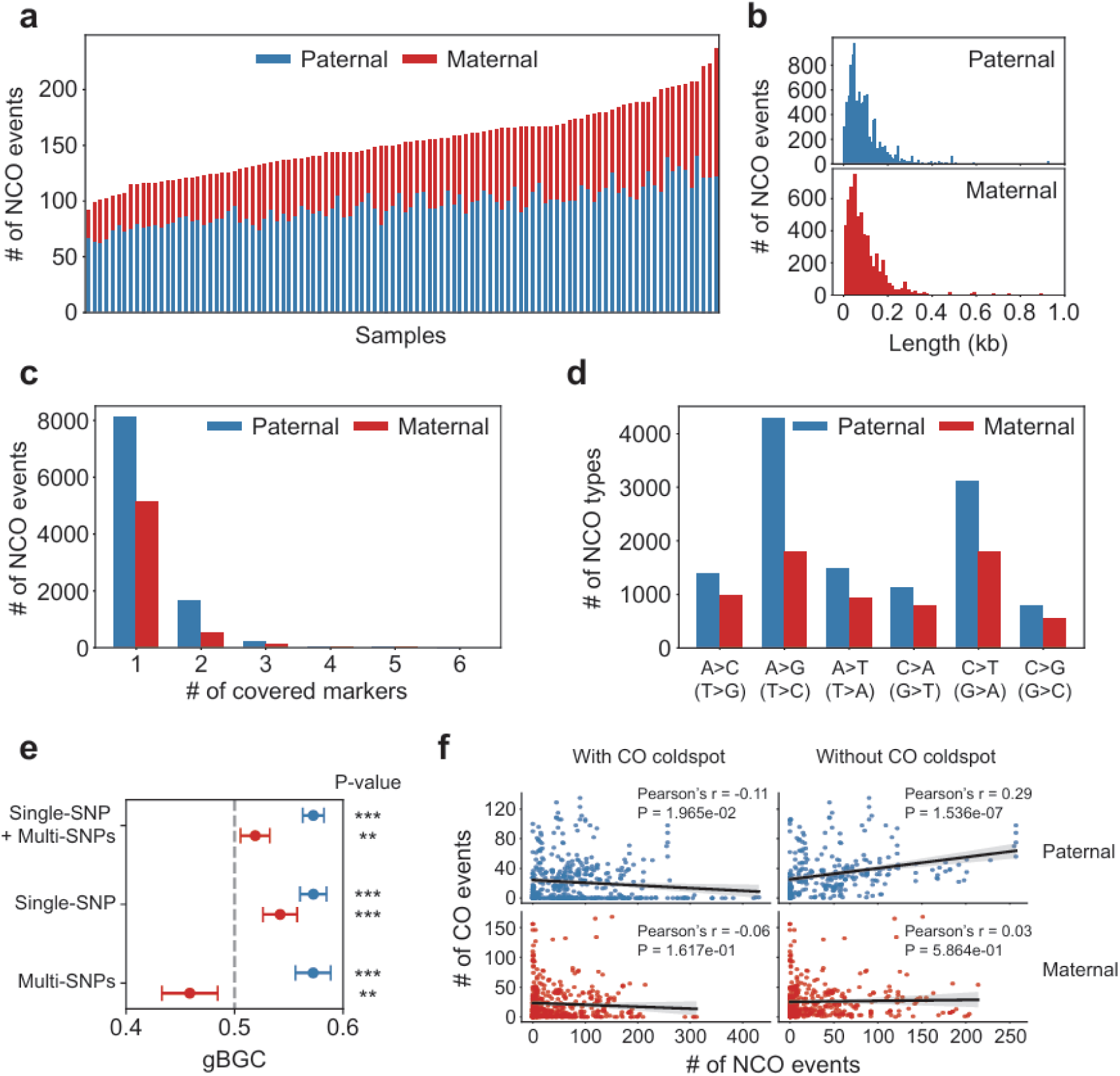
Detection of NCO events. **A**. Numbers of NCO events identified in the paternal and maternal alleles of 104 offspring. **B**. Length distributions of NCO events in paternal (blue barplots, top) and maternal genomes (red barplots, bottom). **C**. Barplots showing the number of NCO events covered by a specific number of SNP markers. **D**. Conversion spectrum of NCO events. A>C includes A>C in the positive strand and T>G in the negative strand, similarly for other types. **E**. NCO conversion bias. NCO events are divided into single-SNP tracts and multi-SNPs tracts according to the numbers of covered markers. X-axis depicts the proportion of A/T > C/G events among ([A/T > C/G] + [C/G > A/T]) events. The p-values and 95% confidence intervals (error bars) were obtained from an exact binomial test. P-values are displayed on the right, with ‘***’ for p-value<0.001 and ‘**’ for 0.001<p-value<0.01. **f**. The relationship between local CO rates and NCO rates in the paternal and maternal genomes, with a sliding window size of 5 Mb. Linear regression was employed to fit the data, with the 95% confidence interval as shaded area. Scatterplots are shown separately for data with and without CO coldspots.

We further analyzed the impacted sequence lengths and conversion types of NCO events. Most NCO events in paternal and maternal genomes involve small genomic regions, with 90.9% being under 200 bp, with the average length of NCO events being 99 bp (**Fig. 5b**). Most NCO events (83.6%) only covered one converted marker (**Fig. 5c**). These NCO events were considered to be single-SNP NCOs, with the rest to be multi-SNP ones. Regarding the conversion types (**Fig. 5d**), we found that in both paternal and maternal genomes, A>G (or T>C) and C>T (G>A) conversions were the most frequent. GC-biased gene conversion (gBGC), common in mammals, favors the fixation of C/G alleles over A/T alleles and mainly occurs in single-SNP conversion tracts^[39]^. Except for the maternal multi-SNP NCOs which showed an AT bias, we discovered GC bias in all other types of NCOs to varying degrees (**Fig. 5e**), with paternal genomes exhibiting higher GC bias. This indicates that gBGC also contributes to NCO events in amphioxus.

We also investigated the relationship between local CO and NCO rates in the paternal and maternal genomes (**Fig. 5f**), with a sliding window size of 5 Mb. We found no or very weak correlation between local CO and NCO rates when including the CO coldspot regions. When excluding CO coldspots, we found a relatively weak positive correlation (Pearson’s r = 0.29, p = 1.536e-7) between local CO and NCO rates in the paternal genome, but no correlation in the maternal genome. This suggests that the local frequencies of CO and NCO events in this species appear to be largely independent.

## Discussion

In this study, we conducted whole genome sequencing for 106 individuals from a two-generation *B. floridae* family to systematically analyze CO and NCO events during male and female meioses. The high heterozygosity of *B. floridae* individuals allowed us to assemble the paternal and maternal genomes into relatively long contigs (average length of ∼5 kb) with only short read data. Based on the offspring read alignments with the parental genomes (assembled contigs), we phased and reconstructed parental haplotypes at the contig level. This method enables phasing of diploid parents without direct gamete sequencing. This approach leverages the high heterozygosity of the amphioxus genome, turning a potential analytical challenge into an advantage. The high genomic heterozygosity also provides a high density of markers for detecting NCO recombination events. If not sequencing gametes, previous genomic research directly detecting recombination events usually needed to sequence pedigrees of three or more generation^[30, 40]^, but our approach only needs two generations. Thus, our strategy offers a new perspective for studying recombination in species with a highly heterozygous genome.

Using this approach, we analyzed CO events in paternal and maternal genomes. The estimated CO recombination rate was higher (paternal: 4.57 cM/Mb, maternal: 4.49 cM/Mb) than that of vertebrates such as mammals (∼0.5 cM/Mb) or birds (∼2 cM/Mb)^[19, 20]^. However, this discrepancy may be partially attributed to the compact genome size of amphioxus relative to vertebrates. In fact, the average number of CO events per haploid chromosome per meiosis was approximately 1.2, close of that of vertebrates^[19]^. Unlike many vertebrates, no obvious sexual dimorphism was found in CO rates in paternal and maternal genomes of this family, but the reason behind this is unclear. Both paternal and maternal genomes have many CO coldspots, likely due to large-scale heterozygous inversions that can suppress CO events^[28]^. Like the high density of SNPs, there are likely many polymorphic heterozygous inversions and other structural variations in the wild amphioxus populations^[29]^. Given that large-scale heterozygous inversions can suppress CO events, there may be large difference in genomic distribution of COs between different individuals, leading to a relatively weak correlation between the directly-measured individual CO recombination map and the historical CO recombination map estimated from population genomic data. Besides, we found that TE and gene densities are positively correlated with local CO rates, though the underlying molecular mechanisms leading to such correlations are unclear. In many vertebrates, the genomic binding profile of PRDM9 (the protein with zinc-finger structures binding specific DNA sequences and triggering histone modifications like H3K4me3/H3K36me3) determines CO hotspots^[41]^. Due to the lack of PRDM9 ChIP-seq data in amphioxus species, we did not perform any PRDM9 related analysis, so its potential role in amphioxus COs remains unknown.

Our analysis of NCO events showed that in amphioxus NCO events occur much more frequently than CO events and have short impact sequence ranges. Like vertebrates, amphioxus also exhibits gBGC, which tends to preserve C/G alleles during conversions^[42, 43]^. Many characteristics of the NCO events in amphioxus are similar to those of vertebrates, suggesting that the mechanisms underlying NCO events may be highly conserved during chordate evolution.

We acknowledge several limitations in our analysis. First, while using bubble contigs as markers for detecting CO recombination events offers higher specificity, bubble contigs are derived from highly heterozygous regions of the genome. This can lead to an uneven genomic distribution of bubble contigs, potentially causing an underestimation of recombination rates in regions with lower heterozygosity. Second, our study sequenced a single family, and thus there are possible biases due to the small sample size. This is especially relevant given that there may be substantial individual-specific heterozygous inversions that can suppress recombination. Consequently, we cannot exclude the possibility that such limitations may mask some differences existing at the population level, such as those linked to sex.

In summary, by sequencing 106 individuals from a *B. floridae* family, we have comprehensively mapped CO and NCO events in amphioxus male and female individuals. This study fills important gaps in our understanding of amphioxus recombination and enhances our knowledge of genetic diversity in amphioxus populations. Our work can facilitate future research of this important model organism in biology.

## Methods

### Sample collection and sequencing

Amphioxus (*B. floridae*) was obtained from a stock maintained by Jr-Kai Yu originating from Tampa, Florida, and the colony and their offspring were raised under previously described conditions^[44]^. 106 individuals from a two-generation family, including two parents and 104 offspring, were adopted in the analysis. DNA was extracted and used to construct paired-end 150 bp sequencing libraries via a shotgun strategy. Whole genome sequencing was performed using the BGISEQ-500 platform. More details can be found in a companion study on amphioxus *de novo* mutation rates^[11]^.

### Genome assembly and determining inherited parental haplotype contigs in offspring

We performed *de novo* haplotype-aware assembly of the parental genomes using Platanus-allee^[10]^ and merged the resultant haplotype contigs into a custom parental reference genome. In Platanus-allee, heterozygous regions are assembled as ‘bubble’ contigs with two allelic sequences, with the longer one named as ‘primary’ and the shorter as ‘secondary’. Homozygous regions with identical sequences in an individual are assembled as ‘non-bubble’ contigs. By aligning offspring sequencing reads to the custom reference genome using BWA^[45]^, we determined which haplotype contig of a specific parent at a divergent locus is inherited in each offspring. Detailed methods can be found in a companion study on amphioxus *de novo* mutation rates^[11]^.

### Identification of CO events

As non-bubble contigs derived from homozygous regions are not informative for recombination analysis, we used only bubble contigs as markers to detect CO events during meiosis in the father and the mother.

The ideal marker bubble contigs should clearly show haplotype inheritance from parents to most offspring. For each bubble contig pair, the ratio of offspring inheriting primary or secondary contigs should be about 1:1. Hence we removed bubble contigs where inheritance couldn’t be determined in over 20% of the offspring or those significantly deviating from a binomial test (P < 0.05), which might introduce bias and inaccurately estimate CO events. We then took the homologous regions from the remaining bubble contig pairs and used nucmer in MUMmer v4.0.0beta2^[46]^ to map them onto the chromosome-level genome assembly^[8]^ to determine their chromosomal positions. We excluded the bubble contigs mapped to multiple chromosomes (possible rearrangement sites) and those unmapped. After filtering, 28,795 bubble contig pairs remained in the paternal genome and 27,673 in the maternal genome for CO detection. We grouped these contigs by chromosome, combined inheritance results from all offspring, and used the R package hapi^[12]^ with maximum parsimony to reconstruct parental chromosomal haplotypes for each parent. By comparing offspring chromosomal haplotypes with parental ones, we identified CO events in the gametes of each offspring.

To identify CO coldspots across the genome, a sliding window approach with a window size of 5 Mb and a step size of 1 Mb was employed. A region was defined as a coldspot if no CO events were detected in any offspring within that region. Successive coldspot regions were subsequently merged to delineate genome-wide coldspot distributions.

### Identification of breakpoints within contigs induced by CO events

To verify and complement our strategy using bubble contigs as markers, we analyzed putative CO-induced breakpoints within bubble contigs. Based on the alignment of offspring sequencing reads to the Platanus-allee parental reference genome, a potential CO breakpoint was identified when an offspring’s reads aligned to both the primary and secondary haplotypes of a bubble contig pair, with the aligned regions being complementary and forming a new haplotype. This indicates a partial sequence exchange between the two original haplotypes. To confirm the breakpoint was from a CO rather than an NCO event, we checked read alignments of all offspring in that locus. If four haplotypes were present in the offspring population, the breakpoint was confirmed as a CO breakpoint.

### Detection of heterozygous inversions

To identify large-scale heterozygous inversions in parents that may suppress CO events, we first used the Platanus-allee^[10]^ assembly and bubble contig phasing results to generate haplotype sequences for each parental chromosome. Then, we aligned these parental haplotypes to a more complete chromosome-level assembly^[8]^ using Minimap2 v2.26-r1175^[47]^ with parameters ‘-a -x asm20 --cs -r2k’. Next, we used SVIM-asm v1.0.3^[48]^ in haploid mode to detect inversions in each haplotype. By comparing variations between the two haplotypes, we identified heterozygous inversions in the parental genomes. Subsequently, the contigs carrying heterozygous inversions were obtained, and the regions with a mapping length of > 500 bp were selected for further analysis and visualization.

### Correlation analysis between CO events and genomic features

To assess the correlation between CO events and genomic features, we scanned the genome using a sliding window approach. Given the sparse distribution of CO events, we used a large window size of 5 Mb with a step size of 1 Mb to reduce random fluctuations from low CO counts. We excluded CO coldspot regions, which are potentially affected by parental heterozygous inversions, from the analysis. Genomic annotations for features like gene density were from Huang et al.^[8]^, and repeat or transposon regions were annotated with RepeatMasker.

### Analysis of NCO events

As non-bubble regions lack markers for NCO detection, we focused on NCO events in bubble regions. By aligning primary and secondary contigs of each bubble contig pair, we identified candidate SNPs. Regions with excessive SNPs may suffer from poor alignment quality, so based on the previously reported heterozygosity (3.2-4.2%) of the amphioxus genome, we selected bubble regions with 1-5 SNPs per 100 bp and used those SNPs inside as NCO markers. We also excluded homologous markers shared by the father and the mother to avoid potential alignment errors across different parental alleles, which could lead to false positive NCOs. Also, as it needs at least three markers to determine if there is an NCO event, we removed bubble contig pairs with fewer than three SNPs.

Based on the alignment data of offspring and parental genomes, we used HaplotypeCaller module of GATK v4.2.0.0^[49]^ with the ‘-ERC GVCF’ parameter to call variants and determine the genotypes of all offspring at the marker sites. An NCO event was identified, when an inherited contig from a bubble contig pair had the same parental phase of markers at both ends but another parental phase in the middle marker(s), with only three haplotypes observed in offspring. Regarding the impacted sequenced range of NCO events, given the inability to obtain the precise length, we used the average of the minimum length (distance between the first and last converted markers, or 1 bp if only one marker was converted) and the maximum length (distance between the nearest markers flanking the converted region) as the approximate length of the NCO events. Although our method cannot distinguish an NCO event from a *de novo* substitution, given the low mutation rate (4.4×10^-9^ per base per generation^[11]^), the level of NCO false positives due to *de novo* substitutions at the heterozygous sites should be very low. For example, of the detected 9,992 paternal NCOs, the number of potential *de novo* substitutions in the surveyed genomic space of 6.71 million (heterozygous markers) × 104 is estimated to be about 1.02 (4.4×10^-9^×6.71×10^6^×104×1/3, with 1/3 assuming equal probabilities of three possible substitutions for a nucleotide), yielding a false positive rate of 0.01%.

## Data Availability

The raw sequence data reported in this paper have been deposited in the Genome Sequence Archive (GSA: CRA027444) that are publicly accessible at https://ngdc.cncb.ac.cn/gsa. Genome assemblies produced by Platanus-allee have been deposited in Science Data Bank (ScienceDB) repository (https://www.scidb.cn/en/anonymous/SmppTUpm). Detailed information of identified CO and NCO events is provided in supplementary tables of this study.

## Code availability

The custom code used for analysis is deposited at https://github.com/tale1209/bf_recom.

## Author contribution

**Lei Tao**: Investigation, Methodology, Formal analysis, Writing - Original draft, Writing - Review & Editing. **Jing Xue**: Investigation, Methodology, Formal analysis, Writing - Review & Editing. **Junwei Cao**: Data Curation, Investigation. **Guang Li**: Conceptualization, Data Curation, Resources, Writing - Review & Editing, Supervision. **Cai Li**: Conceptualization, Resources, Methodology, Writing - Review & Editing, Supervision.

## Conflict of interest

The authors have declared no competing interests.

## Acknowledgments

We thank Dr. Luohao Xu for his assistance in obtaining previous genomic data of amphioxus populations. This work was supported by National Natural Science Foundation of China (32470690), Natural Science Foundation of Fujian Province of China (2022J06004), State Key Laboratory of Biocontrol and Guangdong Provincial Key Laboratory for Aquatic Economic Animals.

## Notes

### Summary of Updates

Update the author list and some text in the manuscript.

